# Genetic inhibition of *PCSK9*, atherogenic lipoprotein concentrations, and calcific aortic valve stenosis

**DOI:** 10.1101/560458

**Authors:** Nicolas Perrot, Donato Moschetta, S. Matthijs Boekholdt, Vincenza Valerio, Andreas Martinsson, Romain Capoulade, Elvira Mass, Marie-Annick Clavel, Nicholas J. Wareham, Christian Dina, Hao Yu Chen, James C. Engert, George Thanassoulis, Patrick Mathieu, Yohan Bossé, Philippe Pibarot, J. Gustav Smith, Marina Camera, Sébastien Thériault, Paolo Poggio, Benoit J. Arsenault

## Abstract

**Background:** Proprotein convertase subtilisin/kexin type 9 (PCSK9) inhibition reduces plasma concentrations of low-density lipoprotein cholesterol (LDL-C), apolipoprotein B (apoB) and lipoprotein(a) [Lp(a)]. Atherogenic lipoprotein levels have been linked with calcific aortic valve stenosis (CAVS). Our objectives were to determine the association between variants in *PCSK9* and lipoprotein-lipid levels, coronary artery disease (CAD) and CAVS, and to evaluate if PCSK9 could be implicated in aortic valve interstitial cells (VICs) calcification.

**Methods:** We built a genetic risk score weight for LDL-C levels (wGRS) using 10 independent *PCSK9* single nucleotide polymorphisms and determined its association with lipoprotein-lipid levels in 9692 participants of the EPIC-Norfolk study. We investigated the association between the wGRS and CAD and CAVS in the UK Biobank, as well as the association between the *PCSK9* R46L variant and CAVS in a meta-analysis of published prospective, population-based studies (Copenhagen studies, 1463 cases/101,620 controls) and unpublished studies (UK Biobank, 1350 cases/349,043 controls, Malmö Diet and Cancer study, 682 cases/5963 controls and EPIC-Norfolk study, 508 cases/20,421 controls). We evaluated PCSK9 expression and localization in explanted aortic valves by capillary Western blot and immunohistochemistry in patients with and without CAVS. Von Kossa staining was used to visualize aortic leaflet calcium deposits. PCSK9 expression under oxidative stress conditions in VICs was assessed.

**Results:** The wGRS was significantly associated with lower LDL-C and apoB (p<0.001), but not with Lp(a). In the UK Biobank, the association of *PCSK9* variants with CAD were positively correlated with their effects on apoB levels. CAVS was less prevalent in carriers of the *PCSK9* R46L variant [odds ratio=0.71 (95% confidence interval, 0.57-0.88), p<0.001]. PCSK9 expression was elevated in the aortic valves of patients with aortic sclerosis and CAVS compared to controls. In calcified leaflets, PCSK9 co-localized with calcium deposits. PCSK9 expression was induced by oxidative stress in VICs.

**Conclusion:** Genetic inhibition of *PCSK9* is associated with lifelong reductions in the levels of non-Lp(a) apoB-containing lipoproteins as well as lower odds of CAD and CAVS. PCSK9 is abundant in fibrotic and calcified aortic leaflets. Oxidative stress increases PCSK9 expression in VICs. These results provide a rationale for performing randomized clinical trials of PCSK9 inhibition in CAVS.

## Introduction

Calcific aortic valve stenosis (CAVS) is the most common form of heart valve diseases and its prevalence is steadily rising in Western societies, affecting almost 3% of the population older than 65 (1). Identification of the risk factors for CAVS could facilitate the development of novel and innovative treatment strategies. To date, the only effective treatments for CAVS are surgical or transcatheter aortic valve replacement and no pharmacological agents have been proven effective for the treatment of CAVS. We and others have shown that drugs which lower low-density lipoprotein cholesterol (LDL-C), such as statins and ezetimibe, do not impede CAVS progression (2–4) or decrease CAVS incidence (5). Coronary artery disease (CAD) and CAVS share many risk factors and pathophysiological mechanisms (6). Whether other cardiovascular drugs could be effective for the treatment of CAVS is unknown.

Proprotein convertase subtilisin/kexin type 9 (PCSK9) is an enzyme secreted by the liver that binds to the LDL receptor (LDLR) and targets it for lysosomal degradation (7). Genetic association studies have shown that natural variations at the *PCSK9* locus (present in 2-3% of the population) are associated with lifelong exposure to low LDL-C levels, and cardiovascular protection (8,9). PCSK9 inhibitors have been shown to substantially lower LDL-C levels in various populations and reduce the risk of adverse cardiovascular outcomes in patients at high cardiovascular risk (10,11).

Lifelong exposure to low LDL-C levels has also been linked to lower aortic valve calcium accumulation and protection against CAVS (12). Recently, investigators of the Copenhagen General Population Study, the Copenhagen City Heart Study, and the Copenhagen Ischemic Heart Disease Study observed that individuals carrying the R46L variant in *PCSK9* are characterized by lower levels of LDL-C, Lipoprotein(a) (Lp[a]) and a lower risk of CAVS (13). However, these results have not been replicated and it is still unclear whether the cardiovascular benefits are due to changes in LDL-C, Lp(a), or both. It also remains unclear whether the protective effect of PCSK9 variants for CAVS is attributable to lower concentrations of apolipoprotein (apo) B-containing lipoproteins, and whether PCSK9 contributes to valvular interstitial cell (VIC) calcification.

The objectives of our study were to identify which parameter(s) of the lipoprotein-lipid profile explained the cardiovascular benefits of genetically-mediated PCSK9 inhibition best and to determine whether variants in *PCSK9* is associated with CAVS in a meta-analysis of four large prospective studies. We also evaluated if PCSK9 was present in fibrotic and calcified aortic valves and if isolated human VICs could produce PCSK9 when under oxidative stress.

## Methods

### Study populations

#### EPIC-Norfolk

The design of the EPIC-Norfolk prospective population study has been published previously (14,15). Participants were identified as having incident CAVS if they were hospitalized with CAVS or if they died with CAVS as an underlying cause. Non-fasting blood samples were drawn into plain and citrate bottles. Blood samples were processed directly at the Department of Clinical Biochemistry, University of Cambridge, or stored at −80°C. Serum levels of total cholesterol, HDL-C, and triglycerides were measured in fresh samples with the RA 1000 (Bayer Diagnostics, Basingstoke, UK). LDL-C levels were calculated using the Friedewald formula. Lp(a) and apoB levels were measured in a subset of the cohort with available stored frozen blood samples. Lp(a) levels were measured with an immunoturbidimetric assay using polyclonal antibodies directed against epitopes in apolipoprotein(a) (Denka Seiken, Coventry, United Kingdom), as previously described (16). This assay has been shown to be apolipoprotein(a) isoform-independent. Serum levels of apoB were measured by rate immunonephelometry (Behring Nephelometer BNII, Marburg, Germany) with calibration traceable to the International Federation of Clinical Chemistry primary standards.

#### UK Biobank

The design of the UK Biobank has also been previously published (17). The present analyses were conducted under UK Biobank data application number 25205. CAVS diagnosis was established from hospital records, using the International Classification of Diseases, 10th revision (ICD10) and Office of Population Censuses and Surveys Classification of Interventions and Procedures (OPCS-4) coding. CAVS was defined as ICD10 code number I35.0 or I35.2. Participants with a history of rheumatic fever or rheumatic heart disease as determined by ICD10 codes I00–I02 and I05–I09 were excluded from the CAVS group. We included all other participants in the control group, except for those with OPCS-4 codes K26 (plastic repair of aortic valve) or K30.2 (revision of plastic repair of aortic valve) or a self-reported diagnosis of CAVS, which were excluded from the analysis. We defined CAD as self-reported myocardial infarction, CAD from ICD or OPCS codes.

#### Malmö Diet and Cancer Study

The Malmö Diet and Cancer Study (MDCS) is a prospective, population-based cohort study from the city of Malmö in southern Sweden. Data collection, sample characteristics, and clinical definitions of prevalent and incident CAVS for MDCS have been described previously (12).

### Genetic association studies in EPIC-Norfolk and UK Biobank

We selected independent (in low linkage disequilibrium) single nucleotide polymorphisms (SNPs; *r*^2^<0.10) at the *PCSK9* locus (within 100Kb of the gene) associated with LDL-C levels at a genome-wide level of significance in the Global Lipids Genetics Consortium (18). This approach yielded 11 SNPs independently associated with LDL-C levels. Of these, 10 were successfully genotyped in the EPIC-Norfolk study. We built a weighted genetic risk score (wGRS) using these 10 SNPs weighted by the effect of each SNP on LDL-C levels. We then assessed the relationship between each of the 10 SNPs individually and the wGRS with plasma lipoprotein-lipid levels [total cholesterol (TC), LDL-C, high-density lipoprotein cholesterol (HDL-C), very-low-density lipoprotein cholesterol (VLDL-C), triglycerides, apoB, apoA-I and Lp(a)] in 9692 participants of the EPIC-Norfolk study using linear regression, adjusted for age and sex. The association between the wGRS and untransformed, natural log-transformed and the square root of lipoprotein-lipid levels was assessed. We also constructed two additional wGRSs using SNPs associated with apoB levels (regardless of their association with LDL-C, 3 SNPs) and another wGRS of SNPs that were associated with LDL-C, but not with apoB (5 SNPs), both scaled to a 1 mmol/L reduction in LDL-C levels. The associations between the wGRSs and prevalent CAD and CAVS in the UK Biobank were tested using logistic regression adjusted for age, sex and the first 10 ancestry-based principal components using R (version 3.5.1). The difference between the two wGRS (one that included 3 SNPs associated with apoB levels and one that included 5 SNPs associated with LDL-C, but not with apoB levels) on CAD and CAVS was assessed using Z-scores.

### Genetic association study of PCSK9 R46L variant and CAVS

We investigated the association between the R46L *PCSK9* variant and CAVS in a meta-analysis of published (Copenhagen General Population Study, Copenhagen City Heart Study, and Copenhagen Ischemic Heart Disease Study, totalling 1463 cases and 101,620 controls (13)) and unpublished (UK Biobank, 1350 cases and 349,043 controls; Malmo Diet and Cancer study, 682 cases and 5963 controls; and EPIC-Norfolk, 508 cases and 20,421 controls) prospective, population-based studies using logistic regression adjusted for age and sex and the first 10 ancestry-based principal components when available. We performed a fixed-effect metaanalysis using the inverse-variance weighted method as implemented in the rmeta package (version 3.0) in R (version 3.5.1).

### Human aortic valve collection

Patients requiring aortic valve replacement due to aortic valve insufficiency and/or CAVS at Centro Cardiologico Monzino IRCCS were prospectively enrolled in this study. Inclusion criteria were elective surgery, age more than 18 years old, ejection fraction >30%, and normal sinus rhythm. Exclusion criteria were prior cardiac surgery, rheumatic heart disease, endocarditis, active malignancy, chronic liver or kidney diseases, calcium regulation disorders (hyperparathyroidism, hyperthyroidism, and hypothyroidism), and chronic or acute inflammatory states (sepsis, autoimmune disease, and inflammatory bowel disease). Aortic valve classification was made by expert cardiac surgeons: 1) normal leaflet structure (i.e. considered as control group for the analyses), 2) increased leaflet thickening (aortic valve sclerosis), and 3) CAVS. Aortic valve specimens were collected from the operating room and processed within 30 minutes. This study was approved by the IRCCS Centro Cardiologico Monzino Institutional Review Board and by the IRCCS Istituto Europeo di Oncologia and Centro Cardiologico Monzino Ethics Committee and informed consent was obtained from all the participants.

### PCSK9 expression in explanted aortic valves

Two μg of whole protein extract from aortic valve leaflets, at a final concentration of 0.5 mg/mL, were used to evaluate PCSK9 expression. Automated capillary Western blot technology (WES, ProteinSimple) was used following manufacturer’s instruction using a 12-230 kDa separation module (ProteinSimple) in combination with anti-PCSK9 (Santa Cruz Biotechnology), anti-GAPDH (Cell Signaling), anti-rabbit and anti-mouse detection modules (ProteinSimple). Quantification was performed by Compass for SW v3.1.7.

### Histological evaluation of PCSK9 expression and aortic leaflet calcification

Tissues were washed three times in PBS 1x and were fixed in 4% paraformaldehyde, dehydrated, included in paraffin, and cut into 5-7 μm slides. Before staining, slides were rehydrated. For PCSK9 staining, slides were incubated in antigen retrieval solution (Target Retrieval Solution Citrate pH 6; Dako cytomation) at 98°C for 20 minutes and then cooled at room temperature for 20 minutes. Slides were treated with 3% H_2_O_2_ to block endogenous peroxidases and then washed 3 times with 1x PBS-Tween (PBST). Blocking solution (1x PBST and 5% BSA) was added and kept for 45 minutes at room temperature. Slides were incubated with primary anti-PCSK9 (Abcam) at 4°C over night. Slides were washed 3 times with 1x PBST and incubated with biotinylated anti-mouse secondary antibody (Vector Laboratories) for 60 minutes at room temperature. Slides were washed 3 times and ABC complex (ABC kit; Vector Laboratories) was added and incubated for 30 minutes at room temperature. Slides were washed 3 times and ImmPACT DAB Peroxidase (HRP) Substrate (Vector Laboratories) was added and let react for 25 seconds. Slides were immediately rinsed in distilled H_2_O and then incubated for 10 seconds with hematoxylin. Slides were dehydrated, mounted with EUKITT (O. Kindler GmbH), and images were taken with AxioScope (Zeiss). Von Kossa staining was carried out with silver nitrate (AgNO_3_) added on top of each section and exposed to ultra-violet illumination at room temperature for 20 minutes. Slides were washed in deionized H_2_O and then incubated in sodium thiosulfate solution 2 g/dL (Sigma-Aldrich) at room temperature for 5 minutes. Slides were washed three times in deionized H_2_O and then incubated for 10 seconds with hematoxylin. Slides were dehydrated, mounted with EUKITT (O. Kindler GmbH), and images were taken with AxioScope (Zeiss).

### Impact of oxidative stress on PCSK9 expression in valvular interstitial cells

Isolation of aortic valve interstitial cells (VIC) was performed using a modified method described by Poggio *et al*. (19). Briefly, aortic leaflets were placed in 2 mg/ml type II collagenase (Worthington Biochemical Corp.) in Advanced Dulbecco’s modified Eagle’s medium (DMEM) containing 10% fetal bovine serum (FBS), L-glutamine 2 mM, 1% Penicillin/Streptomycin solution and incubated at 37°C for 20 min. The loosened endothelial layer was removed and tissues were finely minced and dissociated in type II collagenase (1 mg/ml) for 4 h at 37°C. The resulting VICs were seeded in tissue culture plates in Advanced DMEM media and maintained at 37°C with 5% CO2. All the experiments were performed on cultured cells between their second and fourth passages. To evaluate *PCSK9* expression (Santa Cruz Biotechnologies) under oxidative stress by Western blot, VICs were treated with hydrogen peroxide (H_2_O_2_; 1 mM) in PBS 1x for 1 hour as previously reported (20). VICs were allowed to recover for 48h in Advanced DMEM (Thermo Fisher) then whole protein extract was collected in RIPA buffer 1x supplemented with proteases inhibitor (Thermo Fisher). GAPDH (Cell Signaling) was used as reference. To evaluate VIC calcification potential, VICs were treated with H_2_O_2_ 100 μM every other day for 12 days in Advanced DMEM supplemented with 10% FBS and then calcium quantification was performed by calcium colorimetric assay (BioVision) following manufacturer’s instruction.

### Statistical analysis

For comparison of groups, the results were expressed as mean ± SEM. For continuous data, values were compared between groups with Student’s t-test or ANOVA when two or more than two groups were compared, respectively. Post hoc Dunn’s test was performed when the p value of the ANOVA was < 0.05. Data were analysed with Graph Pad Prism v7.00 and a value of p < 0.05 was considered statistically significant.

## Results

### PCSK9 variants, lipoprotein-lipid levels, coronary artery disease and calcific aortic valve stenosis

The association between the 10 *PCSK9* SNPs as well as the wGRS and natural log-transformed lipoprotein-lipid levels in 9692 participants of the EPIC-Norfolk are presented in Table 1. The wGRS was significantly associated with TC, LDL-C, and apoB (all p<0.001), but not with VLDL-C (although the R46L variant [rs11591147] was associated with lower VLDL-C levels), HDL-C, triglycerides, apoA-I, or Lp(a). Similar associations were found with untransformed and square root-transformed values (data not shown). In the UK Biobank, the wGRS associated with lower LDL-C was associated with lower odds of CAD (OR=0.76 [95% CI=0.70-0.82], p<0.001) per 1 mmol/L reduction in LDL-C (Figure 1). The wGRS associated with lower LDL-C was also associated with a lower CAVS risk (OR=0.55 [95% CI=0.39-0.77], p<0.001) per 1 mmol/L reduction in LDL-C. Of the 10 SNPs strongly associated with LDL-C, only three were significantly associated with apoB levels (p<0.05, Table 1). To further understand which best explained the CAD and CAVS risk reduction associated with *PCSK9* variants, we built two additional risk scores using SNPs associated with apoB levels (regardless of their association with LDL-C) or specifically associated with LDL-C (i.e. not associated with apoB). When scaled to a comparable LDL-C reduction (1 mmol/L), the GRS associated with apoB levels was a stronger predictor of CAD (but not CAVS) risk reduction than the GRS associated with LDL-C levels only (Figure 1). The association between individual SNPs in *PCSK9* and CAD and CAVS are presented in Supplementary Figure 1.

**Figure 1.**
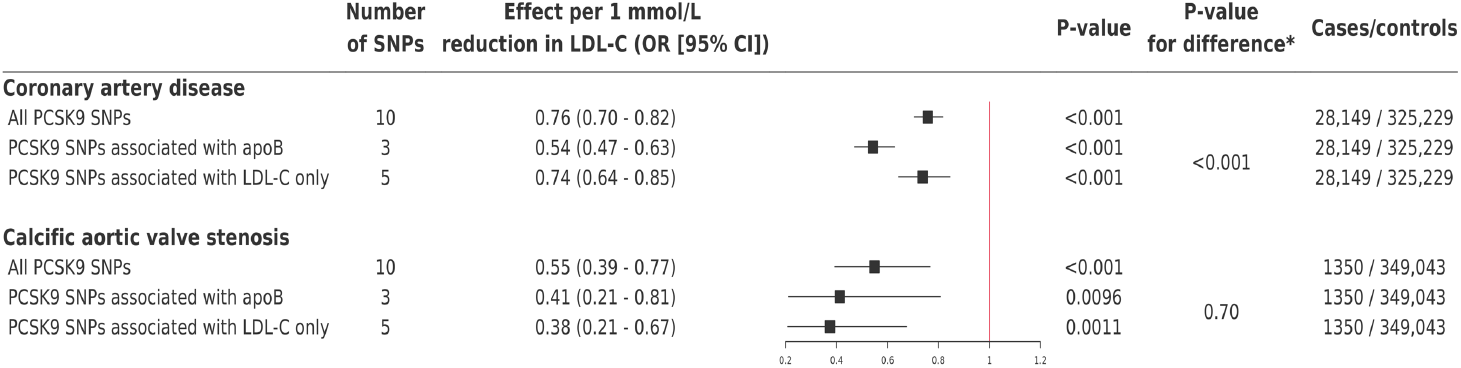
Impact of weighted genetic risk scores of single nucleotide polymorphisms at the *PCSK9* locus associated with low-density cholesterol levels (regardless of their impact on apolipoprotein B levels), apolipoprotein B levels, or LDL-C levels only on coronary artery disease and calcific aortic valve stenosis in the UK Biobank.

**Table 1.** Association of 10 independent, single nucleotide polymorphisms at the *PCSK9* locus with natural log-transformed lipoprotein-lipid levels in 9692 participants of the EPIC-Norfolk study.

|  | TC | P | LDL-C | P | HDL-C | P | VLDL-C | P | TG | P | apoB | P | apoA-I | P | Lp(a) | P |
| --- | --- | --- | --- | --- | --- | --- | --- | --- | --- | --- | --- | --- | --- | --- | --- | --- |
| **rs585131** | -1.2<br>(0.75) | 0.09 | -1.3<br>(0.69) | 0.05 | -0.017<br>(0.0068) | 0.01 | 0.11<br>(0.28) | 0.7 | 3.8<br>(1.3) | 0.00<br>3 | -0.0047<br>(0.0043) | 0.2 | -0.0051<br>(0.0055) | 0.3 | -0.27<br>(0.43) | 0.9 |
| **rs2483205** | -0.15<br>(0.60) | 0.8 | -0.52<br>(0.55) | 0.4 | 0.00013<br>(0.0054) | 1 | 0.37<br>(0.22) | 0.09 | 1.9<br>(1.0) | 0.06 | -0.0010<br>(0.0034) | 0.9 | 0.0046<br>(0.0043) | 0.3 | -0.086<br>(0.34) | 0.9 |
| **rs2479394** | -0.41<br>(0.69) | 0.5 | -0.79<br>(0.63) | 0.2 | 0.012<br>(0.0062) | 0.09 | 0.37<br>(0.25) | 0.2 | -0.32<br>(1.2) | 0.9 | -0.0088<br>(0.0039) | 0.03 | 0.0077<br>(0.0050) | 0.1 | -0.42<br>(0.39) | 0.9 |
| **rs10493176** | -2.2<br>(1.1) | 0.04 | -2.0<br>(0.97) | 0.04 | 0.00082<br>(0.0096) | 1 | -0.18<br>(0.39) | 0.5 | -1.2<br>(1.8) | 0.6 | -0.0060<br>(0.0060) | 0.5 | 0.014<br>(0.0077) | 0.04 | -0.72<br>(0.60) | 0.4 |
| **rs11206510** | -1.7<br>(0.74) | 0.01 | -1.2<br>(0.68) | 0.03 | -0.0099<br>(0.0067) | 0.1 | -0.46<br>(0.27) | 0.05 | -0.65<br>(1.3) | 0.7 | -0.0051<br>(0.0042) | 0.2 | -0.0016<br>(0.0054) | 0.8 | -0.19<br>(0.42) | 1 |
| **rs2479409** | -1.6<br>(0.64) | 0.01 | -1.7<br>(0.58) | 0.003 | 0.0030<br>(0.0057) | 0.6 | 0.12<br>(0.23) | 0.6 | -0.024<br>(1.1) | 0.6 | -0.0024<br>(0.0036) | 0.7 | 0.011<br>(0.0046) | 0.01 | -0.17<br>(0.36) | 0.7 |
| **rs12067569** | -0.33<br>(1.8) | 0.9 | -1.9<br>(1.7) | 0.2 | 0.016<br>(0.017) | 0.4 | 1.6<br>(0.68) | 0.02 | 4.4<br>(3.1) | 0.05 | 0.0057<br>(0.010) | 0.8 | 0.013<br>(0.013) | 0.4 | -0.85<br>(1.0) | 0.4 |
| **rs10888896** | -1.8<br>(0.69) | 0.006 | -2.0<br>(0.63) | <0.001 | -0.0046<br>(0.0062) | 0.5 | 0.18<br>(0.25) | 0.5 | 1.9<br>(1.2) | 0.5 | -0.0039<br>(0.0039) | 0.4 | 1.9e-05<br>(0.0050) | 0.7 | -0.44<br>(0.39) | 0.6 |
| **rs11591147** | -19.8<br>(2.5) | <0.001 | -18.2<br>(2.3) | <0.001 | -0.030<br>(0.023) | 0.2 | -1.6<br>(0.92) | 0.04 | -2.6<br>(4.3) | 0.6 | -0.068<br>(0.014) | <0.001 | -0.0042<br>(0.018) | 0.9 | -0.33<br>(1.4) | 0.6 |
| **rs7552841** | -2.8<br>(0.62) | <0.001 | -2.5<br>(0.57) | <0.001 | -0.0037<br>(0.0056) | 0.5 | -0.32<br>(0.23) | 0.1 | -0.81<br>(1.1) | 0.5 | -0.0093<br>(0.0035) | 0.02 | -0.0015<br>(0.0045) | 0.8 | -0.23<br>(0.35) | 0.7 |
| **wGRS** | -16.9<br>(2.4) | <0.001 | -16.5<br>(2.2) | <0.001 | -0.027<br>(0.022) | 0.2 | -0.41<br>(0.90) | 0.4 | 2.9<br>(4.1) | 0.4 | -0.056<br>(0.014) | <0.001 | 0.018<br>(0.018) | 0.2 | -1.7<br>(1.4) | 0.4 |

Data are presented as beta (standard error). All associations were adjusted for age and sex. TC indicates total cholesterol, LDL-C indicates low-density lipoprotein cholesterol, VLDL-C indicates very-low-density lipoprotein cholesterol, HDL-C indicates high-density lipoprotein cholesterol, TG indicates triglycerides, apoB indicates apolipoprotein B, apoA-I indicates apolipoprotein A-I and Lp(a) indicates Lipoprotein(a).

### Meta-analysis of the association between the PCSK9 R46L variant and calcific aortic valve stenosis

The *PCSK9* R46L variant was associated with lower odds of CAVS in the three new cohorts, although the association was only statistically significant in the UK Biobank. In a meta-analysis of all four cohorts, carriers of the R46L variant had lower odds of CAVS compared to noncarriers (OR = 0.71 [95% CI, 0.57-0.88], p<0.001) (Figure 2).

**Figure 2.**
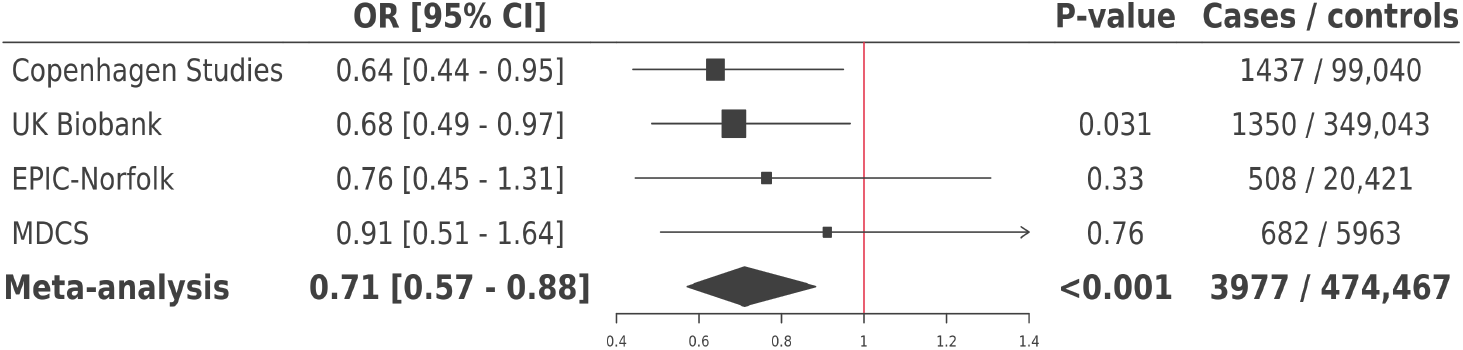
Association of the *PCSK9* R46L variant with calcific aortic valve stenosis in a meta-analysis of four cohorts.

### PCSK9 expression and calcific aortic valve stenosis

We investigated PCSK9 protein expression in explanted aortic valves and found negligible expression of PCSK9 in control aortic leaflets that appeared to be macroscopically normal. Aortic valve whole tissue extract from patients with aortic sclerosis presented significantly higher PCSK9 levels (log_2_-fold change = +28.6±5.1, p=0.008) than controls. Similarly, CAVS patients showed higher PCSK9 levels (log_2_-fold change 39.3±15.2, p=0.02) compared to controls. No statistical differences were observed for sclerotic versus CAVS aortic valves (Figure 3). Immunohistochemical analysis identified PCSK9 in both calcified and non-calcified leaflets. However, PCSK9 was highly abundant in calcified ones (Figure 4 and Supplementary Figure 2). PCSK9 seemed to be expressed by VICs in the fibrosa layer. We observed that microcalcification, as evaluated by Von Kossa staining, co-localized to regions of high PCSK9 expression (Figure 4C right panels). Next, we sought to determine whether CAVS VICs could express PCSK9 under oxidative stress. We treated human isolated VICs with H_2_O_2_ and then evaluated PCSK9 protein levels. PCSK9 expression was induced by oxidative stress (fold change = +2.3±0.4, p=0.02) compared to untreated VICs (Figure 5).

**Figure 3.**
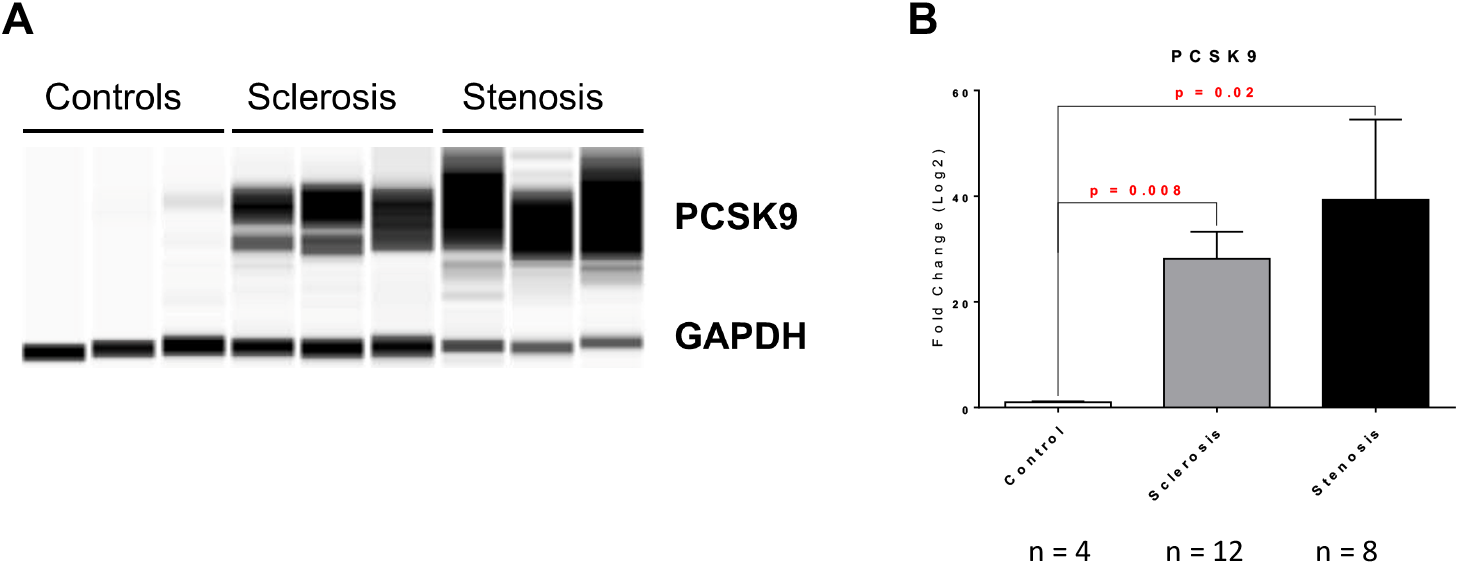
Increased PCSK9 expression in explanted aortic valves. (**A**) Representative image of capillary Western blot showing PCSK9 expression in control (n = 4), sclerotic (n = 12), and stenotic (n = 8) aortic valves. GAPDH has been used for normalization. (**B**) Bar graph showing PCSK9 quantification. The quantification has been plotted as log2-fold change relative to controls.

**Figure 4.**
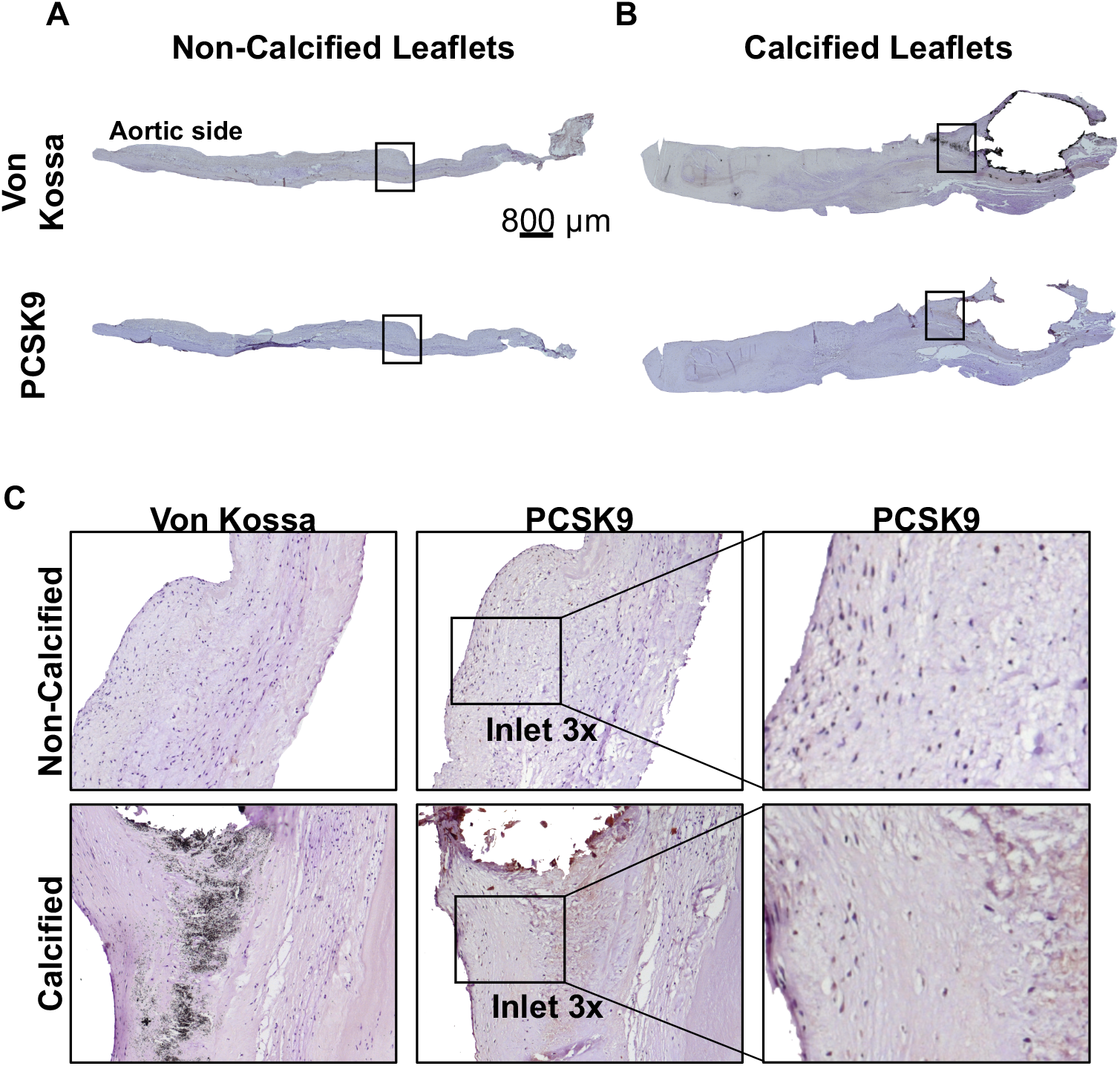
*PCSK9* expression co-localizes with calcium deposits in calcified leaflets. **(A)** Representative non-calcified explanted aortic valve leaflet and **(B)** representative calcified aortic valve leaflet, both stained with Von Kossa to visualize calcium deposits (upper panels) and with anti-PCSK9 (lower panels) 5x. Black boxes indicate the magnification areas (10x). **(C)** 10x magnification of aortic valve leaflets showing co-localization of *PCSK9* expression and calcium deposits (left and middle panels). Right panels indicate inlet digital magnification (3x) of PCSK9 expression.

**Figure 5.**
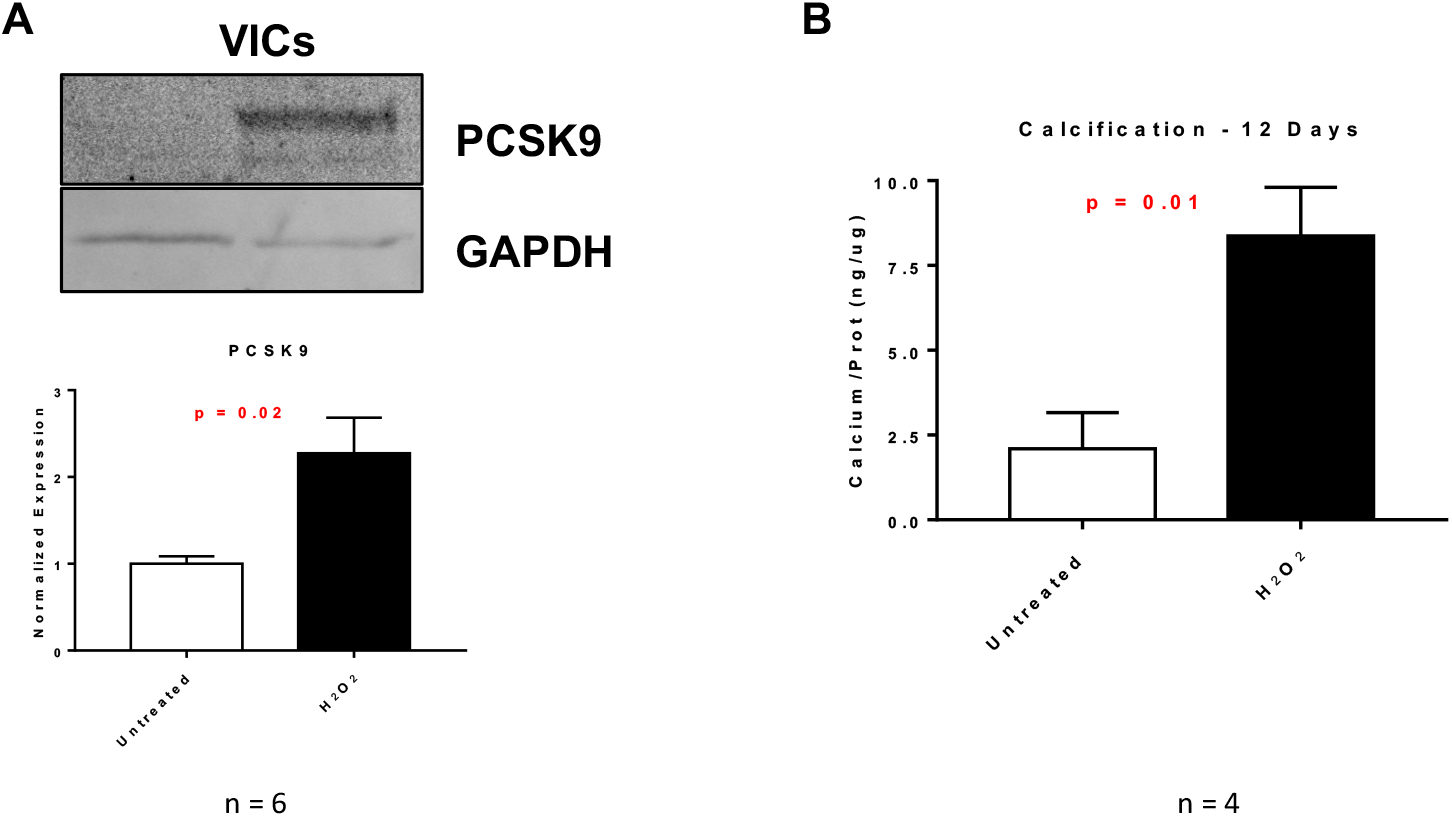
PCSK9 expression is induced by oxidative stress in valve interstitial cells. Representative Western blot showing whole valve interstitial cell (VIC) protein extracts at steady state (untreated) and under oxidative stress (H_2_O_2_ 1 mM for 1 hour and 24 hours of recovery). Bar graph shows PCSK9 protein levels **(A)** normalized to GAPDH expression (n = 6) and calcium **(B)** levels normalized to protein content (n = 4).

## Discussion

Previous genetic association studies observed that carriers of genetic variants in *PCSK9* associated with low LDL-C levels are at lower risk of a broad range of atherosclerotic cardiovascular diseases (21). In this study, we confirmed that variants in *PCSK9* are associated with lower LDL-C levels are not only associated with a lower risk of CAD, but additionally with the risk of CAVS. PCSK9 is present in human aortic valves where it co-localises with calcification areas. Investigating the potential parameters of the lipoprotein-lipid profile that might explain the benefits carrying these variants, we found apoB levels to be a key factor as variants associated with apoB were associated with a higher protection against CAD compared to variants in *PCSK9* associated with LDL-C, but not apoB, levels. In the absence of an association between these variants and Lp(a) levels, we conclude that the cardiovascular benefits of “genetic” PCSK9 inhibition may be attributable to a lifelong lower exposure to non-Lp(a) apoB-containing lipoprotein particles (Figure 6).

**Figure 6.**
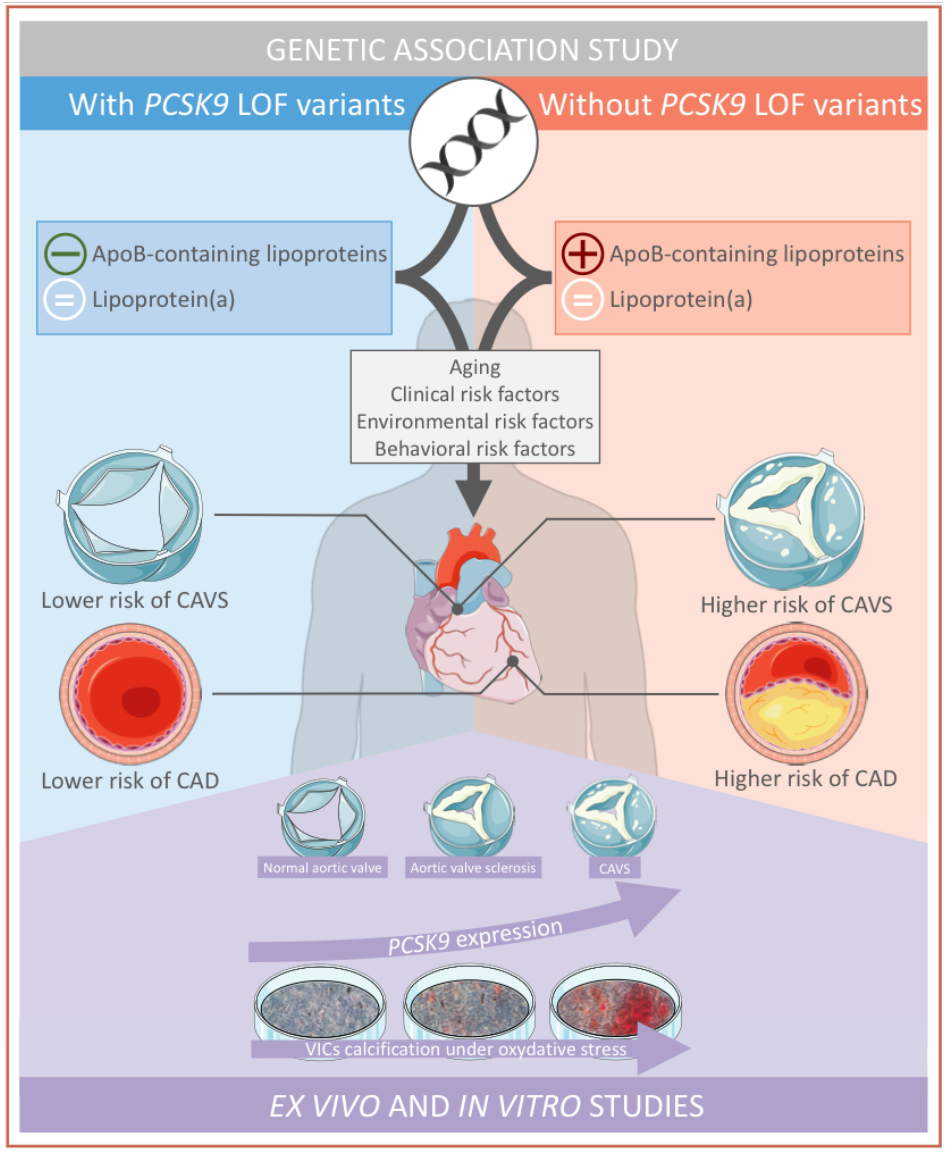
Genetic inhibition of *PCSK9* is associated with lower levels of non-Lipoprotein(a) apolipoprotein B-containing lipoproteins and protection against coronary artery disease and calcific aortic valve stenosis.

Our results are concordant with findings by Ference *et al*. (22) who recently showed that variants in any given gene associated with lower LDL-C track better with cardiovascular protection if these variants simultaneously influence apoB levels, as opposed to those with no effect or a small effect on apoB levels. In the present study, we confirm these findings and expand the investigation of the lipoprotein-lipid profile to apolipoproteins and Lp(a). In a previous report of 2373 individuals included in a nested case-control design of the EPIC-Norfolk study, we have shown that carriers of the *PCSK9* R46L variant had lower levels of nuclear magnetic resonance spectroscopy-measured VLDL and LDL particle concentrations and lower Lp(a) (23). In another study, Sliz *et al*. (24) reported no impact of this variant on VLDL lipid measures. In the current study, we included additional participants and *PCSK9* variants and found no association between SNPs in *PCSK9* and VLDL-C or Lp(a). Although our study population was not as large, our results for Lp(a) are in contrast with those of the Copenhagen investigators who have shown that carriers of the R46L variant (n=1293) have approximately 10% lower Lp(a) levels compared to non-carriers (n=48,324). Consequently, the impact of *PCSK9* variants on Lp(a) is at best modest. Also in support of this premise, results of the FOURIER trial have shown that PCSK9 inhibition with evolocumab provide modest (16%) reduction in Lp(a) levels in patients with high Lp(a) levels (25), suggesting that PCSK9 inhibitors may not exert their cardiovascular benefits through Lp(a) reduction but rather via reductions in other apoB-containing lipoprotein particles. These findings are also supported by the recently published ANITSCHKOW trial that concluded that PCSK9 inhibition with evolocumab was associated with modest reductions in Lp(a) and did not influence arterial wall inflammation assessed by 18F-fluoro-deoxyglucose positron-emission tomography/computed tomography (PET/CT) in patients with hyperlipoproteinemia(a) (26).

In our study, we found a significant association between *PCSK9* variants and CAVS in the UK Biobank. In fact, the impact of a 1 mmol/L reduction in LDL-C levels associated with *PCSK9* variants on CAVS (45% reduction) was nominally superior than its impact on CVD risk (24% reduction). However, given the relatively small number of CAVS cases in the UK Biobank (1350), these results are not definitive and additional studies are needed to confirm these findings and to determine whether the impact of reduced PCSK9 function is independent of the presence of concomitant CAD as approximately half of patients with CAVS also have CAD. Our results confirm those of the Copenhagen cohorts (13) and extend our previous observations that *Pcsk9^−/−^* mice are less likely to develop aortic valve calcification than wild-type mice (27). Results of our previous study also suggest that there is a significant correlation between the amount of PCSK9 produced by VICs and the extent of VIC calcification (27). Our results are also in line with a report from the CHARGE consortium and the MDCS that has shown that a GRS associated with lifelong LDL-C reduction is associated with lower aortic valve calcium accumulation and lower CAVS incidence (12).

Since circulating PCSK9 levels were not measured, we could not determine whether the benefits of carrying *PCSK9* variants go beyond reductions in apoB levels and could be due to lower plasma PCSK9 levels. Nevertheless, our results are in accordance with two other studies of our group showing that circulating PCSK9 levels are associated with bioprosthetic aortic valve degeneration (28,29). Our *ex vivo* data confirmed that PCSK9 is more abundant in the explanted aortic valve from patients with CAVS and aortic sclerosis compared to controls and that PCSK9 is expressed within areas of calcification. *In vitro*, we found that VICs submitted to a pro-calcifying milieu had an increase in PCSK9 levels. Although these experiments were not designed to prove causality of potentially adverse effects of aortic valve PCSK9 expression, we believe that these results are hypothesis generating and further studies will be required to assess the impact of genetic modulation of aortic valve PCSK9 expression on calcification.

Although, our results highlight the potentially causal role of PCSK9 in the etiology of CAVS, the ultimate proof of causality and clinical benefit would be a clinical trial showing that PCSK9 inhibition is linked with a reduction in valvular outcomes. Although, to our knowledge, no such trials are currently planned, an ongoing trial is currently testing the hypothesis that PCSK9 inhibition will reduce aortic valve macrocalcification (measured by computed tomography) and microcalcification (measured by 18F-NaF PET/CT) in patients with mild-to-moderate CAVS (NCT03051360). Our results support the notion that targeting apoB-containing lipoprotein particles is key for optimal CAD risk reduction and potentially CAVS. In the absence of a pharmacological treatment delaying CAVS progression and outcomes, PCSK9 inhibition could provide benefit to patients with CAVS and our results provide a rationale for a large clinical randomized trial to investigate the effect of PCSK9 inhibition in patients with CAVS.

## Acknowledgements

BJA, MAC and ST hold junior scholar awards from the *Fonds de recherche du Québec: Santé* (FRQS). PP holds the Canada Research Chair in Valvular Heart Disease and his research program is supported by a Foundation Scheme Grant from CIHR. RC is supported by a “Connect Talent” research chair from Région Pays de la Loire and Nantes Métropole. PM holds a FRQS Research Chair on the Pathobiology of Calcific Aortic Valve Disease. YB holds a Canada Research Chair in Genomics of Heart and Lung Diseases. GT is supported by R01 HL128550 from the NIH/NHLBI. JGS was supported by grants from the Swedish Heart-Lung Foundation (2016-0134 and 2016-0315), the Swedish Research Council (2017-02554), the European Research Council (ERC-STG-2015-679242), the Crafoord Foundation, Skåne University Hospital, the Scania county, governmental funding of clinical research within the Swedish National Health Service, a generous donation from the Knut and Alice Wallenberg foundation to the Wallenberg Center for Molecular Medicine in Lund, and funding from the Swedish Research Council (Linnaeus grant Dnr 349-2006-237, Strategic Research Area Exodiab Dnr 2009-1039) and Swedish Foundation for Strategic Research (Dnr IRC15-0067) to the Lund University Diabetes Center. EM is supported by the German Research Foundation (Excellence Cluster ImmunoSensation), the Fritz Thyssen Foundation and Daimler and Benz Foundation. The EPIC-Norfolk Study is funded by Cancer Research UK grant number 14136 and the Medical Research Council grant number G1000143.This study was funded by the *Fondation de l’IUCPQ*, the *Fondazione Gigi & Pupa Ferrari ONLUS*, Merck and Pfizer.

## Supplementary material

**Supplementary Figure 1.**
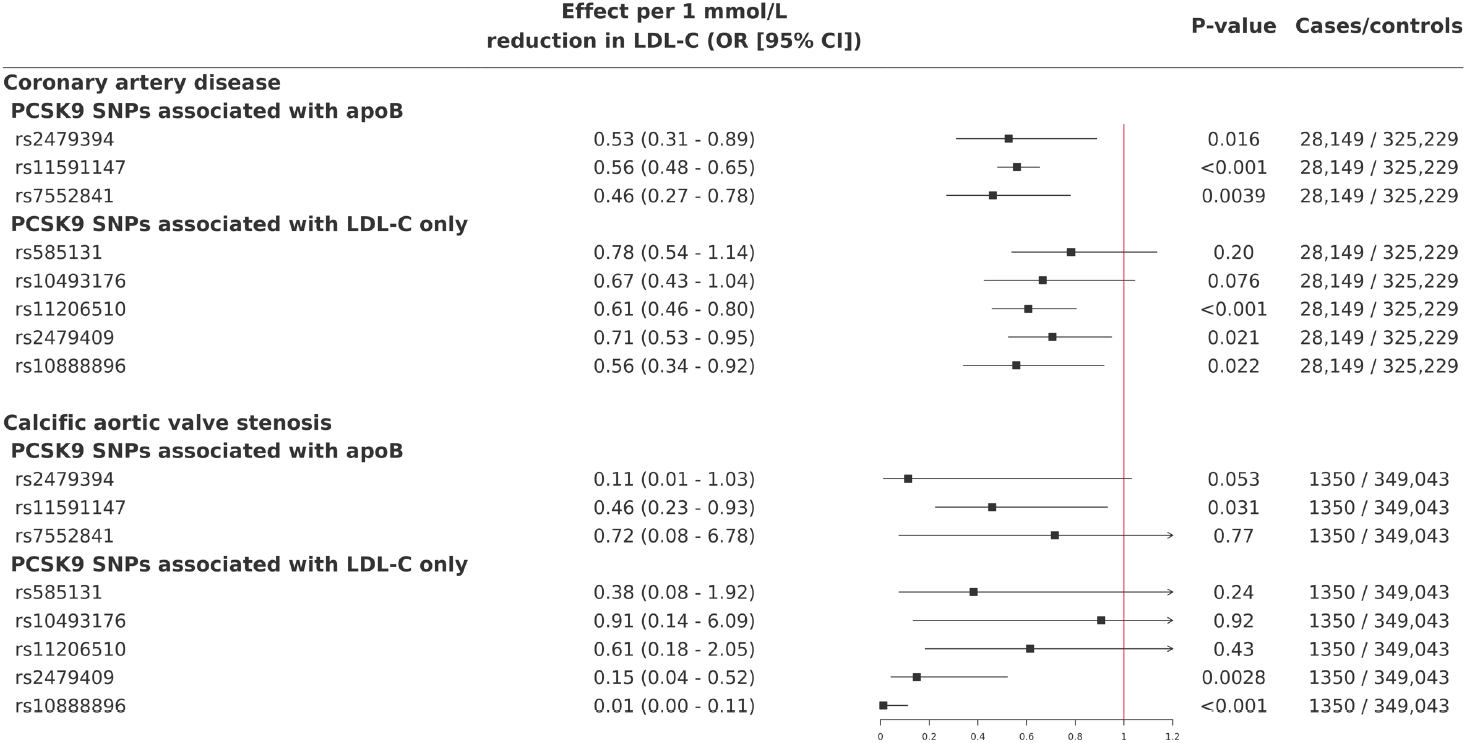
Impact of 10 independent single nucleotide polymorphisms at the *PCSK9* locus associated with low-density cholesterol levels (regardless of their impact on apolipoprotein B levels), apolipoprotein B levels, or LDL-C levels only on coronary artery disease and calcific aortic valve stenosis in the UK Biobank.

**Supplementary Figure 2.**
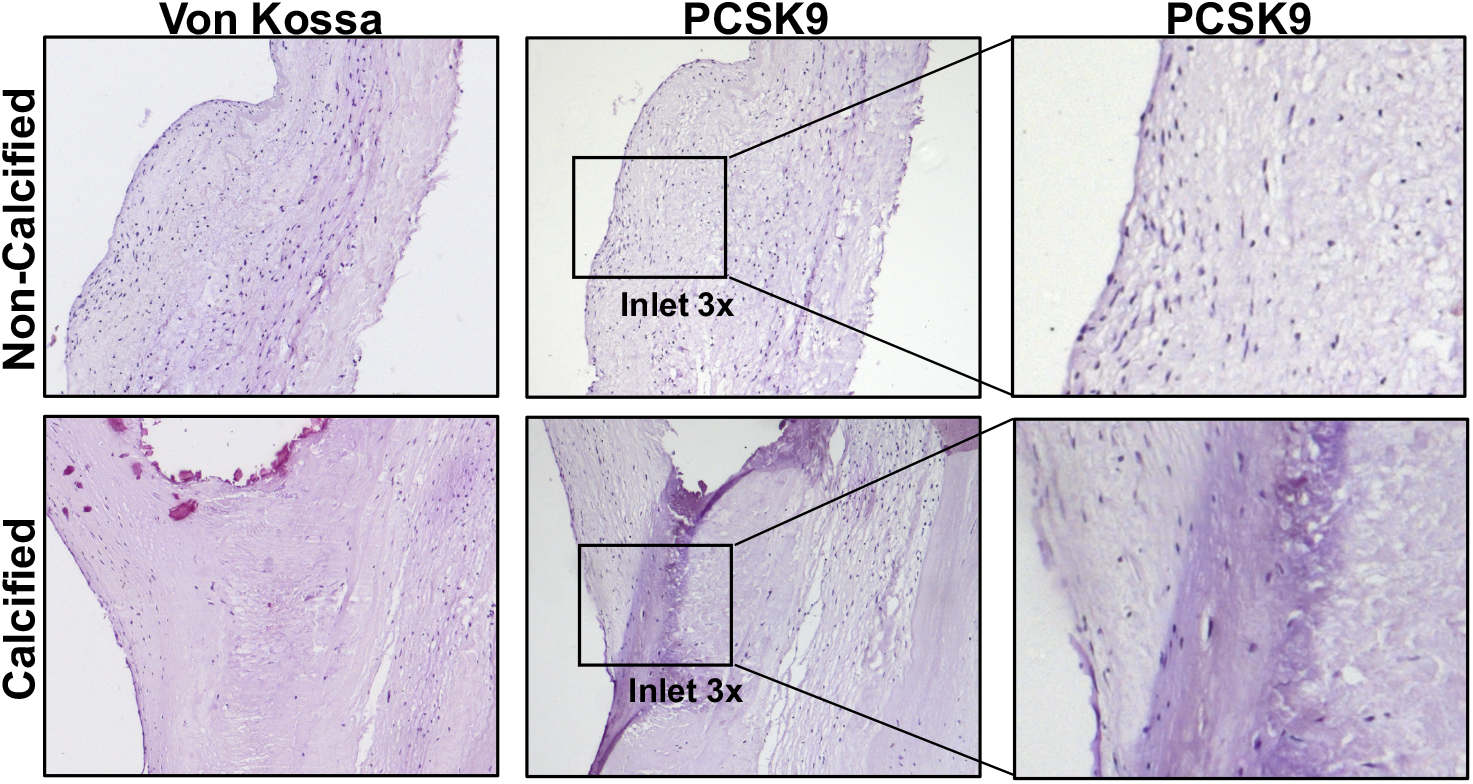
PCSK9 immunohistochemistry and calcium staining negative controls. Representative 10x magnification images of aortic valve leaflets negative control for PCSK9 and calcium staining. Inlet represent 3x digital magnification.

## References

1. Otto CM. Calcific aortic stenosis--time to look more closely at the valve. N Engl J Med 2008;359:1395–8.

2. Rossebo AB, Pedersen TR, Boman K et al. Intensive lipid lowering with simvastatin and ezetimibe in aortic stenosis. N Engl J Med 2008;359:1343–56.

3. Chan KL, Teo K, Dumesnil JG, Ni A, Tam J. Effect of Lipid lowering with rosuvastatin on progression of aortic stenosis: results of the aortic stenosis progression observation: measuring effects of rosuvastatin (ASTRONOMER) trial. Circulation 2010;121:306–14.

4. Cowell SJ, Newby DE, Burton J et al. Aortic valve calcification on computed tomography predicts the severity of aortic stenosis. Clinical radiology 2003;58:712–6.

5. Arsenault BJ, Boekholdt SM, Mora S et al. Impact of high-dose atorvastatin therapy and clinical risk factors on incident aortic valve stenosis in patients with cardiovascular disease (from TNT, IDEAL, and SPARCL). Am J Cardiol 2014;113:1378–82.

6. Perrot N, Boekholdt SM, Mathieu P, Wareham NJ, Khaw K-T, Arsenault BJ. Life’s simple 7 and calcific aortic valve stenosis incidence in apparently healthy men and women. International Journal of Cardiology 2018;269:226–228.

7. Lagace TA, Curtis DE, Garuti R et al. Secreted PCSK9 decreases the number of LDL receptors in hepatocytes and in livers of parabiotic mice. J Clin Invest 2006;116:2995–3005.

8. Cohen JC, Boerwinkle E, Mosley TH, Jr., Hobbs HH. Sequence variations in PCSK9, low LDL, and protection against coronary heart disease. N Engl J Med 2006;354:1264–72.

9. Benn M, Nordestgaard BG, Grande P, Schnohr P, Tybjaerg-Hansen A. PCSK9 R46L, low-density lipoprotein cholesterol levels, and risk of ischemic heart disease: 3 independent studies and meta-analyses. J Am Coll Cardiol 2010;55:2833–42.

10. Sabatine MS, Giugliano RP, Keech AC et al. Evolocumab and Clinical Outcomes in Patients with Cardiovascular Disease. The New England Journal of Medicine 2017;376:1713–1722.

11. Schwartz GG, Steg PG, Szarek M et al. Alirocumab and Cardiovascular Outcomes after Acute Coronary Syndrome. The New England Journal of Medicine 2018;379:2097–2107.

12. Smith JG, Luk K, Schulz CA et al. Association of low-density lipoprotein cholesterol-related genetic variants with aortic valve calcium and incident aortic stenosis. JAMA 2014;312:1764–71.

13. Langsted AG, Nordestgaard BG, Benn M, Tybjærg-Hansen AR, Kamstrup PR. PCSK9 R46L Loss-of-Function Mutation Reduces Lipoprotein(a), LDL Cholesterol, and Risk of Aortic Valve Stenosis. The Journal of Clinical Endocrinology & Metabolism 2016;101:3281–3287.

14. Day N, Oakes S, Luben R et al. EPIC-Norfolk: study design and characteristics of the cohort. European Prospective Investigation of Cancer. Br J Cancer 1999;80 Suppl 1:95-103.

15. Arsenault BJ, Boekholdt SM, Dube MP et al. Lipoprotein(a) levels, genotype, and incident aortic valve stenosis: a prospective mendelian randomization study and replication in a case-control cohort. Circ Cardiovasc Genet 2014;7:304–10.

16. Gurdasani D, Sjouke B, Tsimikas S et al. Lipoprotein(a) and risk of coronary, cerebrovascular, and peripheral artery disease: the EPIC-Norfolk prospective population study. Arterioscler Thromb Vasc Biol 2012;32:3058–65.

17. Sudlow C, Gallacher J, Allen N et al. UK Biobank: An Open Access Resource for Identifying the Causes of a Wide Range of Complex Diseases of Middle and Old Age. PLoS Med 2015;12:e1001779.

18. Willer C, Schmidt E, Sengupta S et al. Discovery and refinement of loci associated with lipid levels. Nature Genetics 2013;45:1274–1283.

19. Poggio P, Branchetti E, Grau JB et al. Osteopontin-CD44v6 interaction mediates calcium deposition via phospho-Akt in valve interstitial cells from patients with noncalcified aortic valve sclerosis. Arteriosclerosis, thrombosis, and vascular biology 2014;34:2086–2094.

20. Branchetti E, Sainger R, Poggio P et al. Antioxidant Enzymes Reduce DNA Damage and Early Activation of Valvular Interstitial Cells in Aortic Valve Sclerosis. Arteriosclerosis, thrombosis, and vascular biology 2013;33:E66-+.

21. Rao AS, Lindholm D, Rivas MA, Knowles JW, Montgomery SB, Ingelsson E. Large-Scale Phenome-Wide Association Study of Variants Demonstrates Protection Against Ischemic Stroke. Circulation Genomic and precision medicine 2018;11:e002162.

22. Ference BA, Kastelein JJP, Ginsberg HN et al. Association of Genetic Variants Related to CETP Inhibitors and Statins With Lipoprotein Levels and Cardiovascular Risk. JAMA 2017;318:947–956.

23. Verbeek R, Boyer M, Boekholdt SM et al. Carriers of the PCSK9 R46L Variant Are Characterized by an Antiatherogenic Lipoprotein Profile Assessed by Nuclear Magnetic Resonance Spectroscopy—Brief Report. Arteriosclerosis, Thrombosis, and Vascular Biology 2017;37:43–48.

24. Sliz EV, Kettunen JO, Holmes MY et al. Metabolomic Consequences of Genetic Inhibition of PCSK9 Compared With Statin Treatment. Circulation 2018;138:2499–2512.

25. O’Donoghue ML, Fazio S, Giugliano RP et al. Lipoprotein(a), PCSK9 Inhibition and Cardiovascular Risk: Insights from the FOURIER Trial. Circulation 2018.

26. Stiekema LCA, Stroes ESG, Verweij SL et al. Persistent arterial wall inflammation in patients with elevated lipoprotein(a) despite strong low-density lipoprotein cholesterol reduction by proprotein convertase subtilisin/kexin type 9 antibody treatment. European heart journal 2018.

27. Poggio P, Songia P, Cavallotti L et al. PCSK9 Involvement in Aortic Valve Calcification. Journal of the American College of Cardiology 2018;72:3225–3227.

28. Salaun E, Mahjoub H, Dahou A et al. Hemodynamic Deterioration of Surgically Implanted Bioprosthetic Aortic Valves. Journal of the American College of Cardiology 2018;72:241–251.

29. Nsaibia MJ, Mahmut A, Mahjoub H et al. Association between plasma lipoprotein levels and bioprosthetic valve structural degeneration. Heart 2016;102:1915.

